# Global patterns of seed germination after ingestion by mammals

**DOI:** 10.1101/549162

**Authors:** Diego A. Torres, John H. Castaño, Jaime A. Carranza-Quiceno

**Affiliations:** Biological Conservation and Biotechnology Research Group, Faculty of Basic Sciences, University of Santa Rosa de Cabal, Colombia

**Keywords:** frugivory, germination patterns, meta-analysis, seed viability

## Abstract

Seed germination is the first step in seedling recruitment. Understanding what factors determine germination success allows some predictions of the effect of climate change or defaunation on the dynamics of plant communities. Mammals play an important role on seed germination through the ingestion of fruit and seeds. However, the populations of many mammal species have been reduced throughout the world, affecting in many ways the dynamics of plant communities. To understand how the loss of mammal populations might impact the dynamics of plant communities first requires us to collect and synthetize all the available evidence on the effects of mammals on seedling recruitment. Here, we used meta-analytical methods to describe the global patterns of the ingestion of seeds by mammals and their effect on seed germination. Our results showed a positive cumulative effect of mammals on seed germination. However, this effect was not the same for all the mammal orders; it varied depending on the plant family and the bioregion. Additionally, increased seed germination was related to rapid germination. These data highlight the important role that some mammals such as Primates and Elephants have on seed germination, and this poses many questions about the mechanisms behind seed germination patterns that will help to guide future research efforts.

## Introduction

Seedling recruitment is the process by which new plants are added to existing populations. This involves three basic steps: seed germination, seedling survival and seedling growth (Eriksson and Ehrlén 2008). The trajectory and success of each of these steps is influenced by complex internal (e.g. seed dormancy and morphology) and external (e.g. resource availability or biotic interactions) factors that act at the individual level through physical, chemical, and ecological mechanisms (Hubbell et al. 2001; Iacona et al. 2010; Willis et al. 2014).

Seed germination, defined as the uptake of water and the elongation of the embryo axis (i.e. the radicle) (Bewley 1997) is the first step in seedling recruitment; and, therefore, it is a crucial determinant of the posterior two steps. The results of seed germination generate patterns further up the individual level, determining the ecology and composition of plant communities (Jiménez-Alfaro et al. 2016). So, understanding the causal factors and mechanisms behind seed germination patterns allows for important applications such as predictions about the effects of climate change (Walck et al. 2011) on defaunation (Wotton and Kelly 2011), or the dynamics of plant communities and ecosystem services, and the development of technologies to improve reforestation (Pinto et al. 2011).

Trophic interactions between plants and vertebrates through fruit and seed ingestion are pervasive factors that have influenced the dynamics of seed germination and seedling recruitment over millions of years (Fleming and Kress 2011). Specifically, seed-ingesting vertebrates affect ingested seeds by changing their spatial position, when they are ingested in one place and defecated in another (Herrera 2002). By changing their immediate environment, for example, dung fertilizing effects (Tjelele et al. 2015) or, by changing their structure after physical and chemical interactions with the gastrointestinal tract (GT); the two most important structural changes are the removal of the pulp (disinhibition effect) and the abrasion and modification of the seed coat (scarification effect) (Traveset et al. 2008).

The effect of vertebrates on seeds is generally dualistic because part of the ingested seeds can be adversely affected, while another part can be positively affected (Genrich et al. 2017). In the case of effects mediated by structural changes, some seeds can be killed during interaction with hard structures of the GT (e.g. teeth or beaks) or by gastric acids (Traveset et al. 2008). Many other seeds are defecated still viable and continue their recruitment process, although from then on they are affected by the structural changes previously suffered. For instance, defecated seeds that are highly scarified by GT can germinate faster than non-scarified seeds (Benítez-Malvido et al. 2014).

Among vertebrates, mammals are a diverse and abundant group of fruit and seed consumers both at high and low latitudes; however, the defaunation processes throughout the world have highly impacted the populations of many of these mammalian species (Ceballos et al. 2017), affecting in many ways the seed germination and recruitment process (Wotton and Kelly 2011). To understand and predict the effects of the loss of mammalian populations on seed germination, recruitment processes, and ecosystem services it is imperative to gather and synthetize all the available evidence of how mammals affect this process. Many published studies have analyzed the effects of individual species or groups of species using similar methods, allowing the use of quantitative research synthesis as methods to explore patterns on a global scale. A first effort was done by Traveset and Verdú (2002), finding that bats increase seed germination more than non-flying mammals, but they did not analyze the effects on the order or family level. More recently, Fuzessy et al. (2016) synthesize the effects of Neotropical primates, finding differences in how feeding guilds affect seed germination percentage and velocity. Finally, Saldaña-Vázquez et al. (in press) synthesized the effects from bats, finding an overall non-effect on seed germination. However, the authors did find significant effects with the analysis at the genus level.

No study has synthetized the available evidence for mammals from all over the world at the same time, and this prevents comparisons at different taxonomic levels and bioregions. Here, we used for the first time meta-analytical methods to synthetize all the available effects of how mammals affect seed germination, the first step in seedling recruitment process. Data synthesis will expose patterns at different taxonomic levels for both mammals and plants, moving forward in the understanding of the functions of mammals and the potential consequences of defaunation. Specifically, we asked 1) what is the global effect that mammals have on the seeds they ingest? 2) Do orders and families of mammals have the same effect? 3) Is the effect the same for different plant families? 4) Is the effect of mammals the same throughout the world? 5) In addition to affecting germination success, do mammals also affect the speed of germination? We expect that patterns of seed germination will open a series of questions and guide future research efforts.

## Materials and methods

### Definitions

Seed germination (SG) is defined as the proportion of seeds that germinate in a seed lot, considering germination as the protrusion of the radicle (in petri dishes or similar germination methods) or as the protrusion above the ground of the hypocotyl (in sowing methods) - this is the so called “visible germination” - because initial stages of seed germination occur at biochemical level (Bewley 1997). Seed viability (SV) is defined as the proportion of seeds in a seed lot stained as viable after an embryonic respiration test with tetrazolium chloride. Mean germination time (MGT) is defined as the mean days from the beginning of the germination test until seed germination occurs in a seed lot. First germination day (FGD) is defined as the days needed for the first seed to germinate in a seed lot.

### Search strategy and study selection

The criteria established for an article to be included in this meta-analysis were as follow: (1) quantification of SG, SV, MGT and FGD, of both a group of ingested seeds (treatment) and a group of non-ingested seeds (control); (2) report the number of seeds of both groups; and (3) seed germination parameters of ingested seeds must come from a mammal. If the study reports more than one species, the parameters should not be combined.

We gathered the studies from *Web of Science* using an advanced search filter created for this study that contained keywords related to seed germination and mammals. After selecting the studies that met the inclusion criteria, the references of those studies were read and other studies meeting inclusion criteria were also selected. This process was done with all selected studies until no more studies meeting inclusion criteria were found. Finally, we did a new search in Google Scholar database using keywords related to seed germination and mammals. Again, the references of all selected studies were read.

### Database

Database contained 17 variables: reference, country, bioregion, latitude, longitude, plant family, plant genus, plant species, mammal order, mammal family, mammal species, effect type, treatment mean, control mean, number of seeds of the treatment, number of seeds of the control, and method of germination. For bioregions we used the classical Wallace’s zoogeographic regions (Wallace 1876), with the update delimitations based on mammals of Kreft and Jetz (2010). The taxonomy of mammals follows “*The Mammal Taxonomy Database*” of the American Society of Mammalogists (Burgin et al. 2018) and plant taxonomy follows “*Plants of the World Online*”, of the Royal Botanic Gardens (http://www.plantsoftheworldonline.org/). When numerical data in a study were reported in figures, these were extracted using the software WebPlotDigitizer (https://automeris.io/WebPlotDigitizer).

### Data analyses

To estimate the effect of ingestion by mammals on SG and SV, we used the natural log of the odd ratio (lnOR), as lnOR = [(I_pos_/C_pos_)/(I_neg_/C_neg_)], where I_pos_ is the number of ingested seeds germinated, I_neg_ is the number of ingested ungerminated seeds, C_pos_ is the number of germinated seeds not ingested, and C_neg_ is the number of ungerminated seeds that were not ingested. We used effect sizes (lnOR) to calculate the cumulative effect sizes by categories (mammal orders, mammal families and plant families). Effect sizes were also arranged in different data sets in order to calculate cumulative effects of mammalian orders by plant families and bioregions.

Since heterogeneity among studies was expected (and, indeed, confirmed by Cochran’s Q and I-square), due to different methods of germination tests and the number and diversity of species included, we used a random-effects model with the DerSimonian-Laird weighting method (DerSimonian and Laird 2015) to calculate the cumulative effect sizes (lnOR_++_). For each effect size we calculated the lower and upper 95% confidence intervals (CI), which were used to estimate the precision of the effect size. When the CIs did not overlap zero we considered the effect size to be statistically significant. We performed the analysis with the software OpenMee (Wallace et al. 2017).

Mammal ingestion effects on MGT and FG were calculated using the natural log of the response ratio (lnRR) as: ln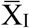 - ln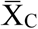, where 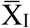 is the mean value of ingested seeds and 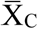 is the mean of non-ingested seeds (Borenstein et al. 2009). Because the standard deviation values were not available in 55% of MGT and 85% FG effect sizes we could not use routine cumulative methods, therefore we used a non-parametric test Kruskal-Wallis in order to test for differences in lnRR of MGT and FGD by mammal orders, followed by a Tukey’s HSD post-hoc test to compare all pairs. To test for relationships between SG and seed germination velocity effects (MGT and FGD) we calculated the Pearson correlation coefficient. We performed the calculation of lnRR on an Excel spreadsheet and we calculated non-parametric ANOVA, post-hoc test, and correlations with the statistical software R, through R commander package (Fox 2005).

## Results

The search resulted in 154 studies from 48 countries in all six bioregions (Fig 1). From those studies we gathered 1116 effect sizes (SG=850, SV=38, MGT=172, FGD=56) from 448 plant species of 104 families and 115 mammal species of 12 orders.

**Figure 1.**
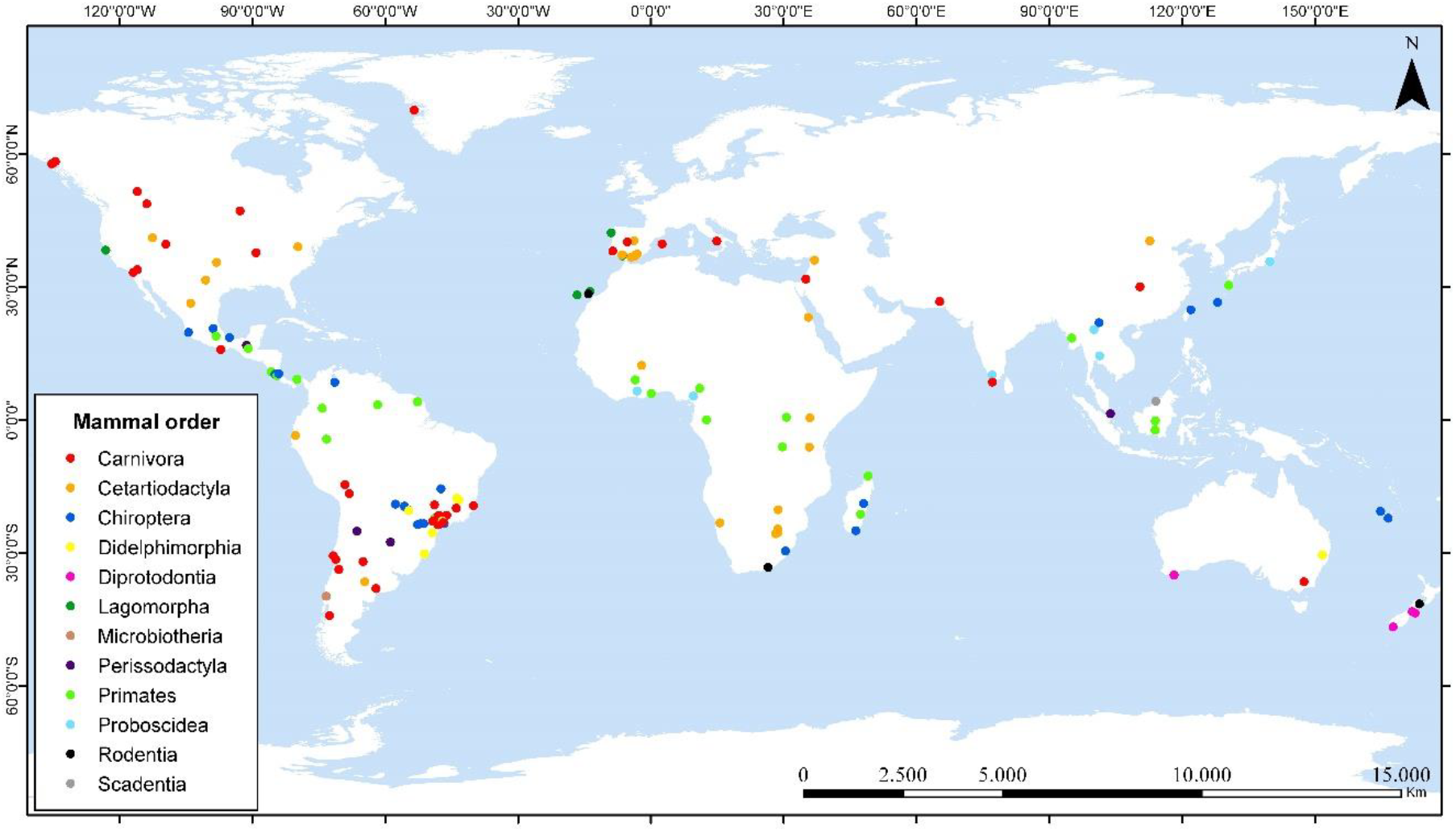
Global distribution of studies included in this meta-analysis.

### Seed germination by mammal orders

The overall effect of mammals on seed germination is positive and significant (lnOR++=0.230; CI=0.139, 0.321). Five orders of mammals (Fig. 2) tend to increase the germination of ingested seeds, but only Proboscidea (lnOR++=1.236; CI=0.632, 1.841), Primates (lnOR++=0.791; CI=0.612, 0.970) and Didelphimorphia (lnOR++=0.383; CI=0.093, 0.673) have a significant effect. Only one effect size was available for Microbiotheria, which also tend to increase the germination of seeds (lnOR=0.40; CI=0.006, 0.795). The other six orders tend to decrease seed germination (Fig. 2), but only diprotodonts (lnOR++=−0.725; CI=−1.406, −0.048) and rodents (lnOR++=−1.078; CI=−1.931, −0.225) decrease significantly.

**Figure 2.**
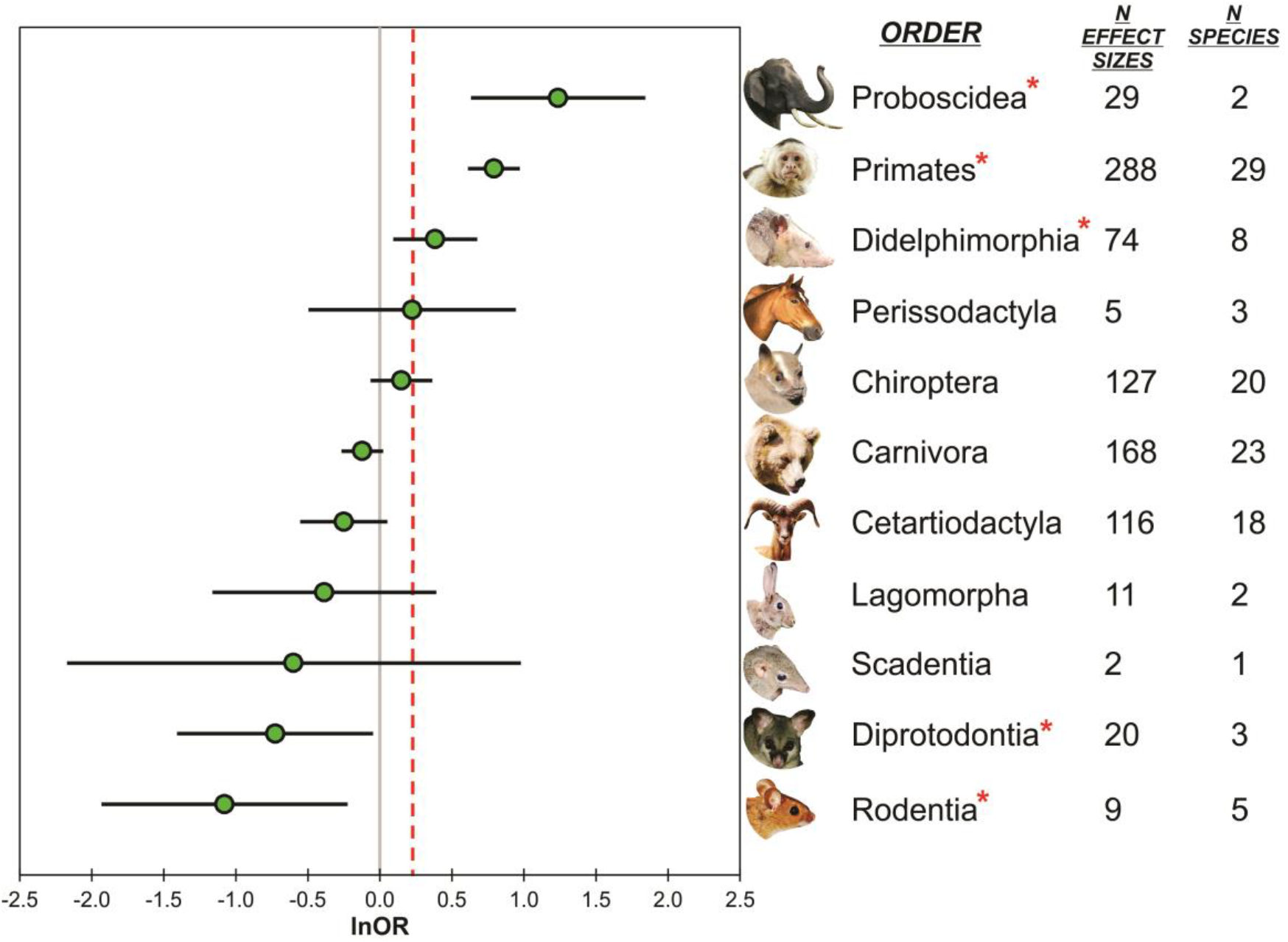
Seed germination by mammal orders. Cumulative effect sizes are reported with their 95% confidence intervals. Effect is significant if the confidence intervals do not overlap zero.

### Seed germination by mammal families

Effect sizes from 30 mammal families were available, with 11 families significantly affecting the germination of seeds positively and 5 families affecting negatively. Families of Australian marsupials Macropodidae and Potoroidae showed the highest effects, however they were calculated based only on one effect size. After Australian marsupials, primate families, except Cebidae (Fig. 3A), as well as Elephantidae were the families that most increased germination.

**Figure 3.**
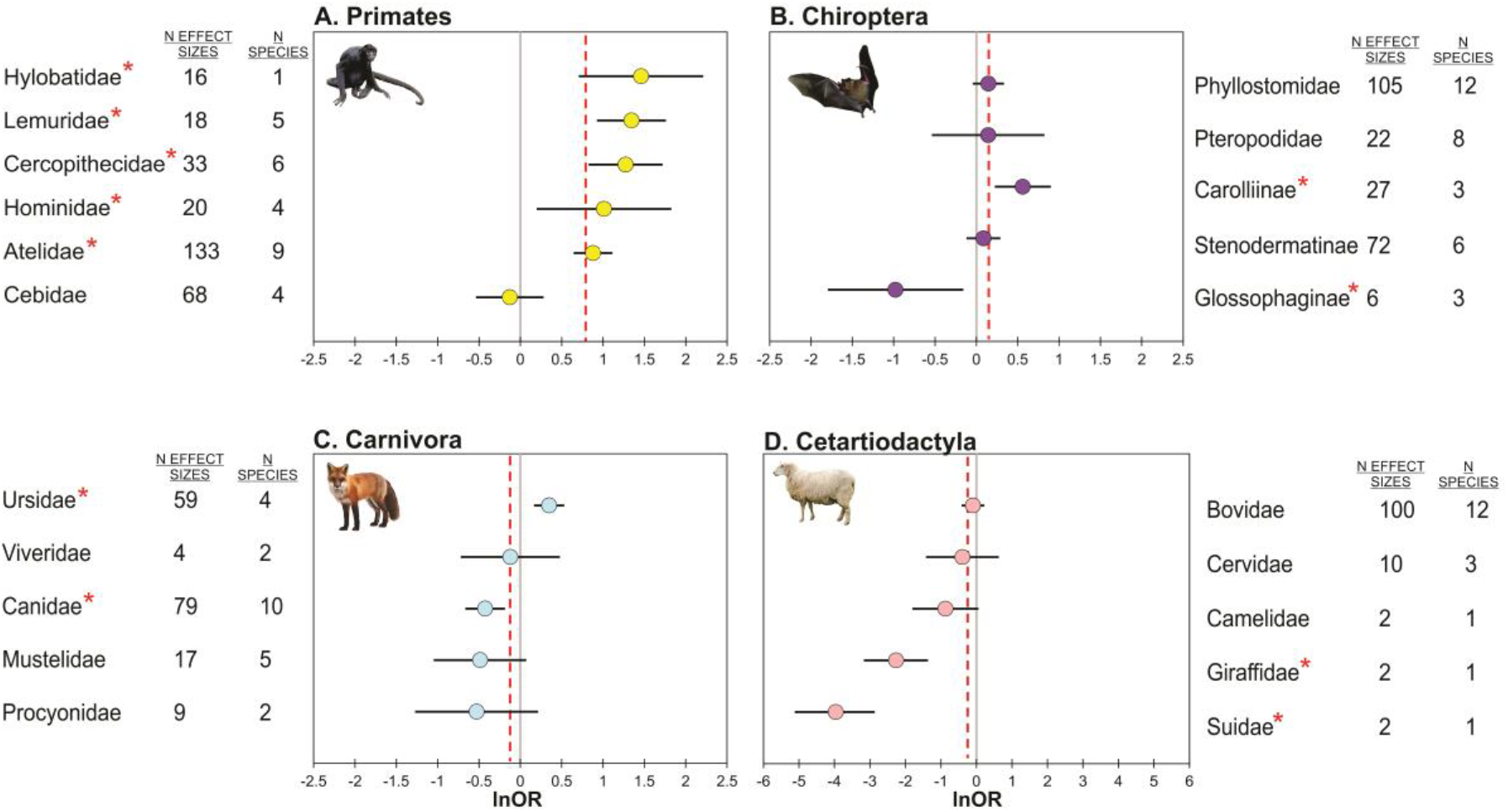
Seed germination by mammal families. Cumulative effect sizes are reported with their 95% confidence intervals. Effect is significant if the confidence intervals do not overlap zero.

The two analyzed bat families, Phyllostomidae and Pteropodidae, had a similar non-significant cumulative effect (Fig. 3B), but analysis of subfamilies of Phyllostomidae showed that frugivorous bats of Carolliinae increased the seed germination significantly (lnOR++=0.560; CI=0.223, 0.898), while nectarivorous bats of Glossophaginae decreased it (lnOR++=−0.978; CI=−1.796, −0.158).

Among carnivores (Fig. 3C), only bears (Ursidae) had a positive effect on seed germination (lnOR++=0.347; CI=0.166, 0.530). The other four families of carnivores analyzed tended to decrease the germination, particularly Canidae, which showed a significant effect (lnOR++=−0.426; CI=−0.665, - 0.186). All families in Cetartiodactyla tended to decrease germination, but only Giraffidae (lnOR++=−2.263; CI=−3.170, - 1.370) and Suidae (lnOR++=−3.963; CI=−5.115, - 2.864) had a significant effect (Fig. 3D).

### Seed germination by plant families

Effects from 103 plant families were available. Seeds from 61 families tended to increase the germination, and 21 were significantly affected. Another 41 families tended to decrease seed germination after mammalian ingestion, with 21 families significantly affected. Only seeds from family Anisophylleaceae (*Poga oleosa*) showed an lnOR=0, when ingested by *Loxodonta cyclotis* (Elephantidae), however this was calculated based only on one effect size. Families with cumulative effects with more than ten individual effect sizes are illustrated in Figure 4.

**Figure 4.**
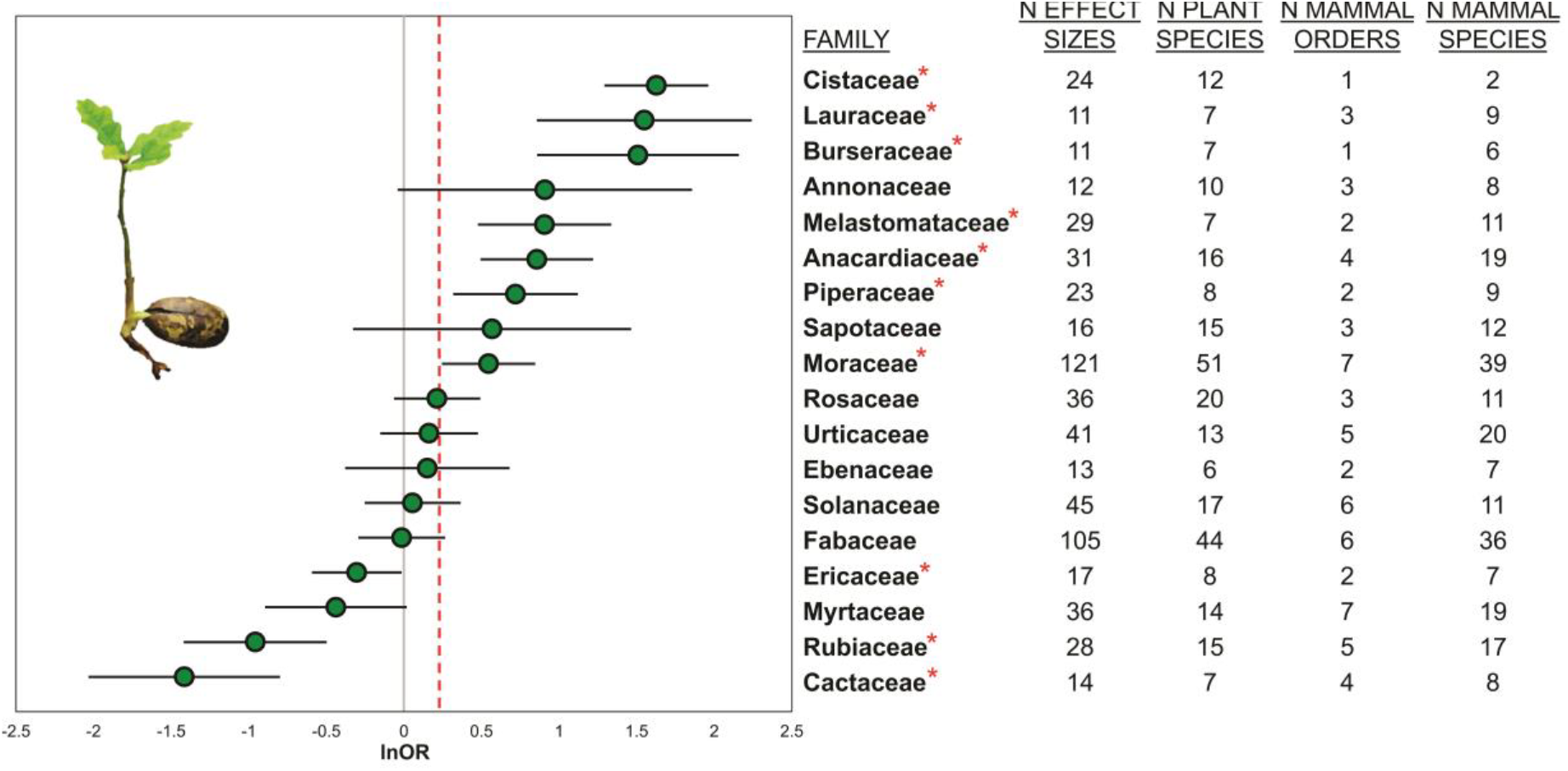
Seed germination by plant families. Cumulative effect sizes are reported with their 95% confidence intervals. Effect is significant if the confidence intervals do not overlap zero.

### Seed germination by plant families and mammal orders

In order to analyze the effect of particular mammal orders on plant families, only families were selected with more than ten effect sizes and more than one mammal order (Fig. 5). Primates were the group that most increased germination in five families, and Cetartiodactyla and Carnivora were the orders that most decrease germination in three families. Some orders that showed an overall non-significant effect in seed germination (Fig. 2) when analyzed by plant families showed significant differences; Carnivora decreased Annonaceae and Myrtaceae, but increased Lauraceae; Cetartiodactyla decreased Cactaceae and Fabaceae; Lagomorpha decreased Rubiaceae and Ericaceae; and Chiroptera increased seeds from Piperaceae.

**Figure 5.**
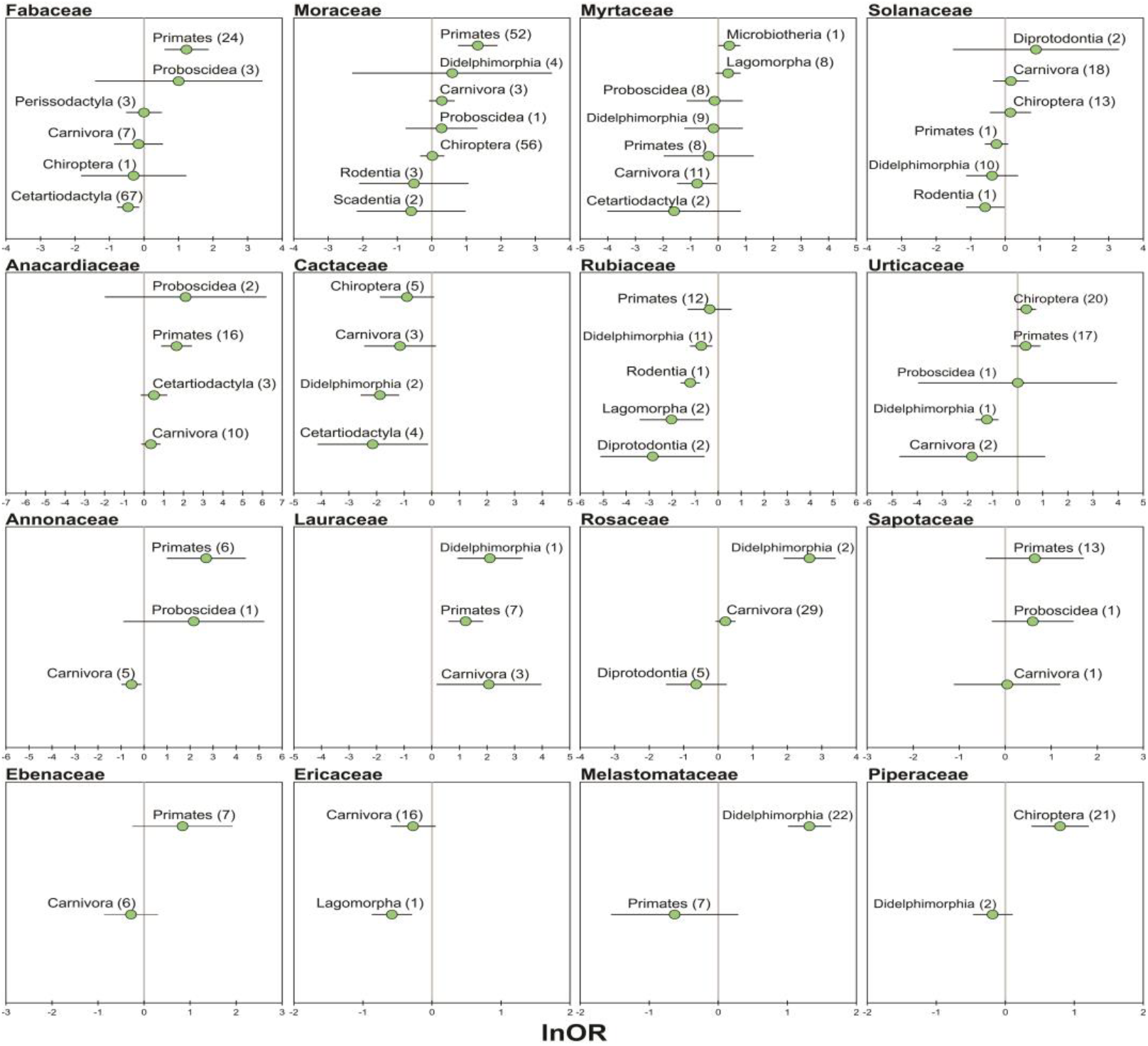
Seed germination by plant families and mammal orders. Cumulative effect sizes are reported with their 95% confidence intervals. Effect is significant if the confidence intervals do not overlap zero.

Again when analyzed by families of plants some mammal orders with particular tendencies in overall effects (Fig. 2) sharply changed their tendency (Fig. 5). Primates tended to decrease seed germination from Rubiaceae, Myrtaceae and Solanaceae. Didelphimorphia significantly decrease seed germination from families Rubiaceae, Urticaceae and Cactaceae and tended to decrease non-significantly seed germination from Myrtaceae, Solanaceae and Piperaceae. Order Carnivora tended to increase seed germination in Moraceae, Solanaceae and Anacardiaceae. Lagomorpha tended to increase germination in family Anacardiaceae and Diprotodontia in family Solanaceae.

### Seed germination by bioregions

Mammal orders had different effects in each bioregion (Fig. 6). The Africa bioregion showed the highest overall positive cumulative effect of mammals on seed germination (lnOR++=0.674; CI=0.407, 0.941), followed by Palaearctic (lnOR++=0.650; CI=0.331, 0.969) and Neotropics (lnOR++=0.318; CI=0.197, 0.440). The Australian bioregion showed the lowest overall negative cumulative effect (lnOR++=−0.489; CI=−0.92, 0.440), followed very closely by the Neartic (lnOR++=−0.449; CI=−0687, −209). The Oriental region showed non-significant overall cumulative effect (lnOR++=0.058; CI=−0.358, 0.474). Some orders such as Cetartiodactyla and Carnivora changed from positive effects to negative effects or vice versa in different bioregions. Most analyzed orders in the Australian and Nearctic showed negative effects, while in other bioregions like Neotropics most analyzed orders showed positive effects.

**Figure 6.**
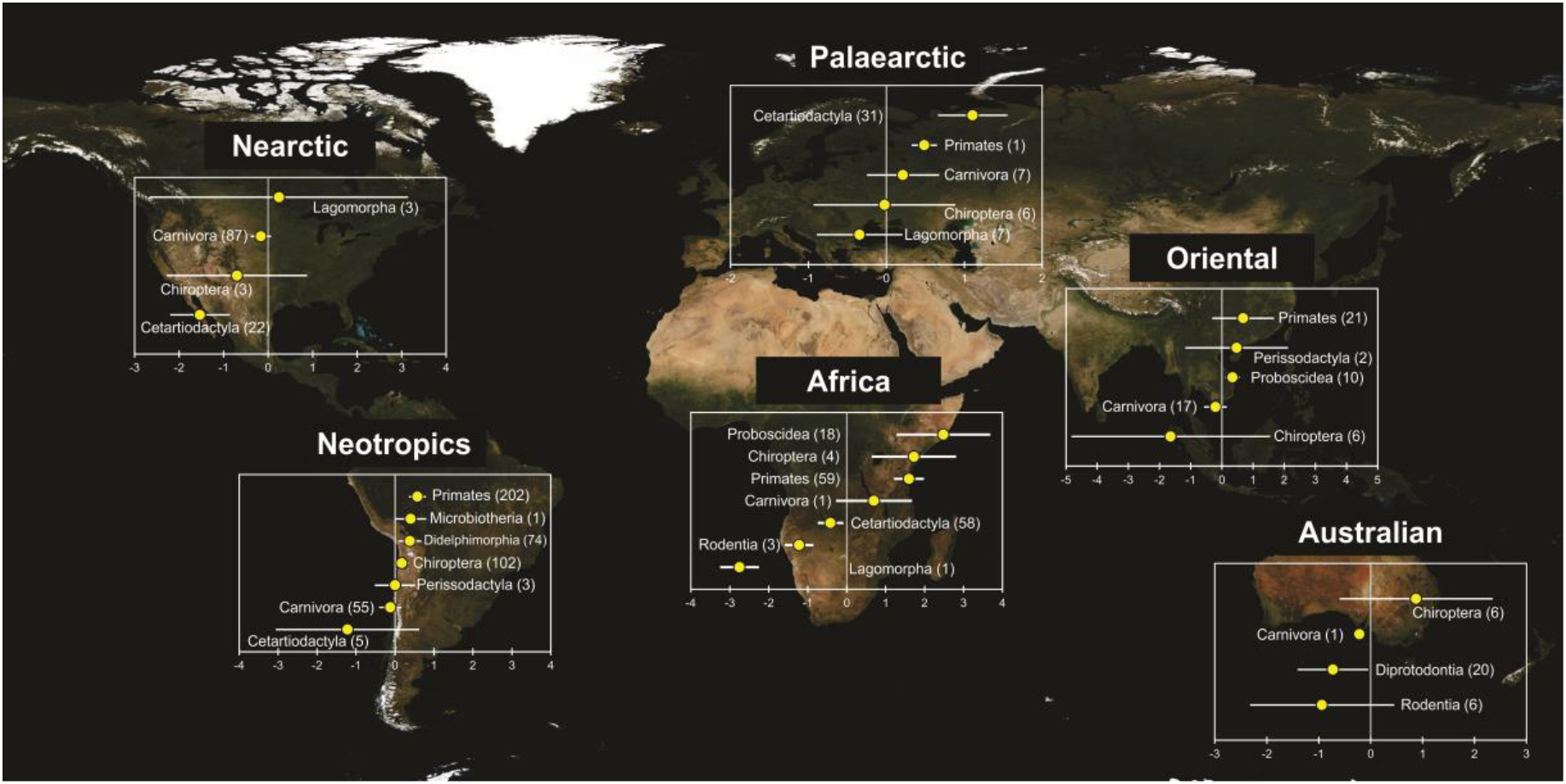
Seed germination of mammal orders at the six Wallace’s bioregions, modified by Kreft and Jetz (2010). Cumulative effect sizes are reported with their 95% confidence intervals. Effect is significant if the confidence intervals do not overlap zero. x-axis represent the lnOR++ of seed germination.

### Seed germination time

Mammal orders had significant different effects on MGT, with Proboscidea showing the shortest times and Lagomorpha the longest times (Fig. 7A). Effects on FGD were also different among mammal orders, but not statistically significant (Fig. 7B). Both MGT and FGD were negative related to SG (Fig. 7C and D), indicating that high germinability is related to fast germination.

**Figure 7.**
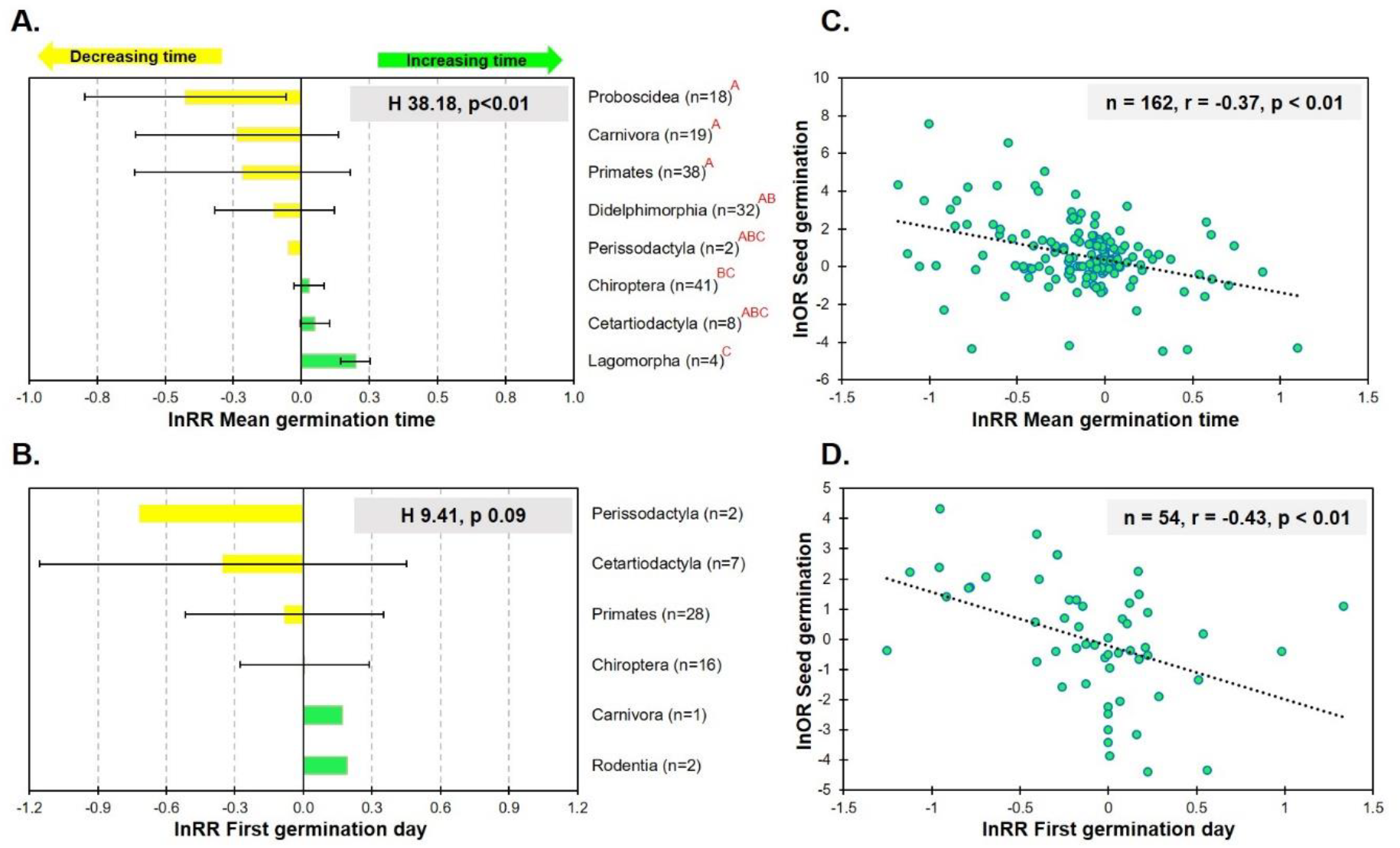
Mean ± SD of the lnRR of Mean germination time (A) and First germination day (B) among mammal orders and the negative correlations with the lnOR of Seed germination (C and D). Different letters (small letters in red) indicate differences after a Tukey’s HSD post-hoc test.

### Seed viability

Effect sizes in seed viability were only available for five mammalian orders. Effect tendencies were similar to seed germination effects, with significant negative effects for the orders Rodentia (only *Atlantoxerus getulus* x *Rubia fruticosa;* lnOR=−2.995; CI=−3.963, - 2.017), Cetartiodactyla (n=16; lnOR++=−1.491; CI=−2.079, −0.901), Lagomorpha (n=8; lnOR++=−1.142; CI=−2.120, −0.161) and Carnivora (n=12; lnOR++=−0778; CI=−1.187, - 0.368), and Diprotodontia (only *Trichosurus vulpecula* x *Crataegus monogyna*) with positive effect, but not significant (lnOR=0.006; CI=−1.624, 1.640).

## Discussion

The synthesis of the available evidence on how mammals affect seed germination shows diverse patterns, which is to be expected by the high number of mammal and plant species interacting. These patterns are composed by a whole range of cumulative effects from negative to positive, changing at different taxonomic levels and from one bioregion to another. The synthetized evidence highlights the important role that some mammals groups have on recruitment processes, turning them into targets for conservation agendas, and also raises new questions and guides future research efforts.

The positive cumulative effect that mammals have on seed germination supports the view that mammals are an important group of plant mutualists in terrestrial ecosystems (Fleming and Sosa 1994), at least as increasers of seed germination. However, not all mammal species have the same effect. The meta-analysis by Traveset and Verdú (2002) finds a differential effect between flying and non-flying mammals, both with significant positive effects but with flying mammals mostly increasing seed germination. That pattern highly contrasts with our results because bats tended to increase seed germination but not significantly, as found by (Saldaña-Vázquez et al.), and non-flying mammals had both negative and positive effects. These contrasting results are probably due to the low number of effects accumulated and the diversity of species in Traveset and Verdú (2002). This also points out the temporality of patterns emerging from synthetic studies, which undoubtedly change as more evidence becomes available (Koricheva and Gurevitch 2013).

When analyzed at the family and subfamily level the patterns of seed germination remained similar to the patterns observed at the order level, suggesting that phylogenetic affiliation may be an important factor shaping the effect on seed germination. However, it is interesting to note that Cebidae (Primates), Ursidae (Carnivora) and Glossophaginae (Chiroptera) sharply contrasted with the pattern at the level of order, therefore on those taxa could be clues about the characteristics that determine the effect of seed ingestion on seed germination. For instance, the cumulative effect of Cebidae was calculated on evidence from *Saguinus mystax, Leontocebus fuscicollis, Callithrix penicillata* and *Cebus capucinus*, all of them omnivorous primates, feeding primarily on insects, gums and fruits (Porter 2001; Mckinney 2011; Vilela and Del-Claro 2011). This result is similar to a previous meta-analysis of Neotropical Primates, which found that primates from the insectivore-frugivore feeding guild increased seed germination less than when compared to frugivore and folivore-frugivore guilds (Fuzessy et al. 2016).

As in Cebidae, the evidence in Glossophaginae also suggested that diet type (and the related physiology and anatomy) is an important determinant of the effect on seed germination. The cumulative negative effects in this subfamily were calculated based on evidence from the nectarivorous and pollinivorous bats *Glossophaga longirostris, G. commissarisi*, and *Leptonycteris yerbabuenae*. In the last species the authors suggest that the decrease of seed germination was caused by the acids of the GT, which killed embryos (Rojas-Martinez et al. 2015). However, the reduction of germination in nectarivorous bats could also be the opposite, because these bats have simple GTs and when they ingest fruits and seeds the digestion is less efficient (Kelm et al. 2008), probably reducing the scarification of seeds and generating the decreased pattern of seed germination that the authors observed.

As mentioned above, the Ursidae significantly increased seed germination, sharply contrasting with the other Carnivora families Viverridae, Canidae, Mustelidae and Procyonidae. The length of the GT of carnivores is usually short compared to other terrestrial mammals (McGrosky et al. 2016). This reduces the time of GT-seed interaction, what in turn decreases scarification and seed germination. Additionally, in carnivores the GT length has a positive relationship to body mass (McGrosky et al. 2016), so it is not surprising that seeds consumed by bears weighing hundreds of kilograms (Jones et al. 2009), produce enough scarification for seed germination to occur at high percentages, as we observed.

When the effect of seed ingestion on seed germination was analyzed by grouping seeds by plant families, diverse patterns emerge, with some families highly favored when consumed by mammals, while other families were highly disfavored. Furthermore, most plant families were not equally affected by all mammal orders, and some plant families showed contrasting patterns of seed germination depending on the mammal taxon that consumed the seeds. This favoring of seed germination has been attributed to the coevolution of some groups of vertebrates with some groups of plants (Charles-Dominique 1986; Cypher and Cypher 1999). However, evidence supporting this hypothesis is still scarce (Saldaña-Vázquez et al.; Traveset 1998). Whether or not seed germination is favored after seed ingestion is the result of coevolutionary processes, will be ultimately determined by the interaction between the seed traits specific to the plant family and the traits of the GT of the mammal order.

Patterns of seed germination changed from one bioregion to another, with some mammals such as elephants and primates preserving their positive effects throughout the bioregions where they occur, whereas other mammals such as bats or carnivores are less conservative on their positive effects. These complex patterns are expected because plant and mammal species composition, and the pairs of interacting species, are not the same in each bioregion. Indeed, only 3 plants and 9 mammals were shared among two or more bioregions. Even cosmopolitan mammals have different effects in each bioregion. For instance, goats feeding on seeds from Fabaceae trees have neutral effects on seed germination in the Nearctic (Kneuper et al. 2003) and the Neotropics (Ortega-Baes et al. 2002), while in Africa they have negative, positive and neutral effects (Miller 1995; Tjelele et al. 2012, 2015).

Although it is generally believed that germination enhancement is beneficial for a plant, it is not always the case. As early stated by Traveset (1998), if the germination enhancement is beneficial and thus adaptive, it will depend on the context in which the germination occurs. Therefore if an increase in the fitness of the plant is not evident, it is not possible to determine whether the increase in germination is advantageous. In addition to the increase in germination, we also found a relationship of this increase to the increase in germination velocity. Again, germinating faster does not necessarily represent a benefit for the plant (Leverett et al. 2018), since faster germination can place the seed, and the seedling, in an unfavorable situation both in terms of intra or interspecific competition and environmental conditions (Orrock and Christopher 2010). Thus, the benefit of germinating and germinating faster always depend on context.

Most studies of seed germination are based on visible germination, which is the elongation of the embryonic axis and the breaking of the seed coat. However, the initial steps of germination are not visible and occur at biochemical level (Bewley 1997). The effect of seed ingestion on SV, quantified using biochemical analysis, agreed with the patterns that emerge from SG, and also suggested that low germination in Rodentia, Cetartiodactyla and Lagomorpha is caused by the death of the embryo and not a lack of scarification. These three mammal orders have long GTs, with large fermentation chambers (Hofmann 1989; Lovegrove 2010), which increases the time of GT-seed interaction and the probabilities of negative change in the structure of the seed coat and killing of the embryo.

The patterns found in this study open many questions and help to guide future research efforts. It will be very useful to understand the proximate mechanisms behind patterns of seed germination after being ingested by mammals, particularly those related to scarification and how it interacts with the morphology and physiology of seeds and GTs and the ecological characteristics of the mammals with dietary specialization. In order to understand the role of scarification it would be valuable to use biochemical methods that determine when germination is absent if this is caused by an excess of scarification and death of the embryo or by a lack of enough scarification. Finally, a deep understanding of germination patterns will also require disentangling the role of phylogeny and coevolution between mammals and plants.

